# Fanconi Anemia FANCM/FNCM-1 and FANCD2/FCD-2 are required for maintaining histone methylation levels and interact with the histone demethylase LSD1/SPR-5 in *C. elegans*

**DOI:** 10.1101/265876

**Authors:** Hyun-Min Kim, Sara E. Beese-Sims, Monica P. Colaiácovo

**Affiliations:** Department of Genetics, Harvard Medical School, Boston, MA 02115; School of Pharmaceutical Science and Technology, Tianjin University, Tianjin 300072, China

**Keywords:** LSD1/SPR-5, FANCM/FNCM-1, FCD2/FCD-2, Histone demethylation and DNA repair, germline

## Abstract

The histone demethylase LSD1 was originally discovered as removing methyl groups from di- and monomethylated histone H3 lysine 4 (H3K4me2/1), and several studies suggest it plays roles in meiosis as well as epigenetic sterility given that in its absence there is evidence of a progressive accumulation of H3K4me2 through generations. In addition to transgenerational sterility, growing evidence for the importance of histone methylation in the regulation of DNA damage repair has attracted more attention to the field in recent years. However, we are still far from understanding the mechanisms by which histone methylation is involved in DNA damage repair and only a few studies have been focused on the roles of histone demethylases in germline maintenance. Here, we show that the histone demethylase LSD1/CeSPR-5 is interacting with the Fanconi Anemia (FA) protein FANCM/CeFNCM-1 based on biochemical, cytological and genetic analyses. LSD1/CeSPR-5 is required for replication stress-induced S-phase checkpoint activation and its absence suppresses the embryonic lethality and larval arrest observed in *fncm-1* mutants. FANCM/CeFNCM-1 re-localizes upon hydroxyurea exposure and co-localizes with FANCD2/CeFCD-2 and LSD1/CeSPR-5 suggesting coordination between this histone demethylase and FA components to resolve replication stress. Surprisingly, the FA pathway is required for H3K4me2 maintenance regardless of the presence of replication stress. Our study reveals a connection between Fanconi Anemia and epigenetic maintenance, therefore providing new mechanistic insight into the regulation of histone methylation in DNA repair.

## INTRODUCTION

Most eukaryotes package their DNA around histones and form nucleosomes to compact the genome. A nucleosome is the basic subunit of chromatin composed of ~147bp of DNA wrapped around a protein octamer comprised of two molecules each of four highly conserved core histones: H2A, H2B, H3, and H4. Core histones can be replaced by various histone variants, each of which is associated with dedicated functions such as packaging the genome, gene regulation, DNA repair, and meiotic recombination (TALBERT AND HENIKOFF 2010). Both the N- and C-terminal tails of core histones are subjected to various types of post-translational modifications including acetylation, methylation, SUMOylation, phosphorylation, ubiquitination, ADPribosylation, and biotinylation.

Histone demethylases have been linked to a wide range of human carcinomas (PEDERSEN AND HELIN 2010). Dynamic histone methylation patterns influence DNA double-strand break (DSB) formation and DNA repair, meiotic crossover events, and transcription levels (ZHANG AND REINBERG 2001; CLEMENT AND DE MASSY 2017). However, the mechanisms by which histone modifying enzymes coordinate their efforts to signal for the desired outcome are not well understood, and even less is known about the role of histone demethylases in promoting germline maintenance.

The mammalian histone demethylase LSD1 was originally discovered as an H3K4me2/1 specific demethylase (SHI *et al.* 2004). Studies in flies and fission yeast revealed increased sterility in the absence of LSD1, however, the underlying mechanism of function by which LSD1 promotes fertility remained elusive (DI STEFANO *et al.* 2007; LAN *et al.* 2007; RUDOLPH *et al.* 2007). *C. elegans* studies suggested it plays a role in meiosis and LSD1/CeSPR-5 mutant analysis revealed a progressive sterility accompanied by a progressive accumulation of H3K4meon a subset of genes, including spermatogenesis genes (KATZ *et al.* 2009). In addition to transgenerational sterility, our previous studies discovered that this histone demethylase is important for double-strand break repair (DSBR) as well as p53-dependent germ cell apoptosis in the *C. elegans* germline (NOTTKE *et al.* 2011), linking H3K4me2 modulation via SPR-5 to proper repair of meiotic DSBs for the first time. Other studies supporting the importance of histone methylation in the regulation of DNA damage repair have attracted more attention to the field in recent years (HUANG *et al.* 2007; KATZ *et al.* 2009; BLACK *et al.* 2010; MOSAMMAPARAST *et al.* 2013; PENG *et al.* 2015). However, the mechanisms by which histone demethylation is involved in DNA damage repair remain unclear and only a few studies have been focused on its roles in germline maintenance.

A growing body of work supports a role for components from the Fanconi Anemia (FA) pathway in DNA replication fork arrest in addition to inter-strand crosslink (ICL) repair (ADAMO *et al.* 2010; SCHLACHER *et al.* 2012; RAGHUNANDAN *et al.* 2015; LACHAUD *et al.* 2016). Here, we show that the histone demethylase LSD1/CeSPR-5 interacts with the Fanconi Anemia (FA) FANCM/CeFNCM-1 protein based on biochemical, cytological and genetic analyses. LSD1/CeSPR-5 is required for hydroxyurea (HU)-induced S-phase DNA damage checkpoint activation and its absence suppresses the embryonic lethality and larval arrest displayed in *fncm-1* mutants. We show that FANCM/CeFNCM-1 re-localizes upon HU exposure and co-localizes with FANCD2/CeFCD-2 and LSD1/CeSPR-5. We also show that the potential helicase/translocase domain of FANCM/CeFNCM-1 is necessary for recruiting FANCD2/CeFCD-2 to the site of replication arrest. Surprisingly, the FA pathway is required for H3K4me2 maintenance regardless of the presence of replication stress. Our study reveals a link between Fanconi Anemia and epigenetic maintenance, therefore providing new insights into the functions of the Fanconi Anemia pathway and the regulation of histone methylation in DNA repair.

## MATERIALS AND METHODS

### Strains and alleles

*C. elegans* strains were cultured at 20°C under standard conditions as described in Brenner (BRENNER 1974). The N2 Bristol strain was used as the wild-type background. The following mutations and chromosome rearrangements were used: LGI: *fncm-1(tm3148), spr-5(by101), hT2[bli-4(e937) let-?(q782) qIs48]*(I; III); LGIV: *spo-11(ok79), nT1 [unc-?(n754) let-?(m435)] (IV; V), fcd-2 (tm1298), opIs406[fan-1p∷fan-1∷GFP∷let-858 3′UTR + unc-119(+)](KRATZ et al. 2010).*

### Transgenic animals

The following set of transgenic worms was generated with CRISPR-Cas9 technology as described in (KIM AND COLAIACOVO 2014; KIM AND COLAIACOVO 2015c; NORRIS *et al.* 2015). In brief, the conserved potential helicase motifs were mutated in FNCM-1 *(fncm-1(rj43*[S154Q]) and *fncm-1(rj44*[M24TN E248Q K250D]) animals as described in (KIM AND COLAIACOVO 2014; KIM AND COLAIACOVO 2015c; KIM AND COLAIACOVO 2016). The FNCM-1 tagged animal *(rj45[fncm-1∷GFP∷3xFLAG])* was created with a few modifications of the CRISPR-Cas toolkit as described in (NORRIS *et al.* 2015). The SPR-5 tagged animal (*rj18*[*spr-5*::GFP::HA + loxP *unc-119(+)* loxP]) I; *unc-119*(ed3) III) was generated as described in (DICKINSON *et al.* 2013). All transgenic lines were outcrossed with wild type between 4 to 6 times.

### Analysis of FNCM-1 protein conservation and motifs

FNCM-1 homology searches and alignments were performed using T-COFFEE (http://tcoffee.crg.cat/) (DI TOMMASO *et al.* 2011). Pfam and Prosite (release 20.70) were used for zinc-finger motif predictions (SONNHAMMER *et al.* 1997).

### Plasmids

sgRNAs targeting *fncm-1* were created as described in (NORRIS *et al.* 2015; KIM AND COLAIACOVO 2016). In brief, the top and bottom strands of the sgRNA targeting oligonucleotides (5µl of 200 µM each) were mixed and annealed to generate double-stranded DNA which then replaced the *Bam*HI and *Not*I fragment in an empty sgRNA expression vector (pHKMC1, Addgene #67720) using Gibson assembly (NORRIS *et al.* 2015; KIM AND COLAIACOVO 2016).

To build the *fncm-1*∷GFP∷FLAG donor plasmid, genomic DNA containing up and downstream ~1kb homology arms were PCR amplified and cloned into the multi cloning site of the pUC18 plasmid along with GFP and FLAG tags synthesized by IDT. To build the *spr-*5∷GFP∷HA donor vector, *spr-5* genomic DNA containing up and downstream ~1kb homology arms together with GFP∷HA + loxP *unc-119*(+) loxP were cloned into the ZeroBlunt Topo vector as described in (DICKINSON *et al.* 2013).

### DNA microinjection

Plasmid DNA was microinjected into the germline as described in (FRIEDLAND *et al.* 2013; TZUR *et al.* 2013; KIM AND COLAIACOVO 2016). Injection solutions were prepared to contain 5 ng/µl of pCFJ90 (P*myo-2*∷mCherry; Addgene), which was used as the co-injection marker, 50-100 ng/µl of the sgRNA vector, 50 ng/µl of the *Peft-3*Cas9-SV40 NLS*tbb-2* 3′UTR and 50-100 ng/µl of the donor vector.

### Monitoring S-phase progression in the germline

Nuclei in the *C. elegans* germline are positioned in a temporal-spatial manner and both mitotic as well as meiotic S-phase progression can be monitored at the distal tip (JARAMILLO-LAMBERT *et al.* 2007). To monitor S-phase progression in the germline, ~ 200pmol/µl Cyanine 3-dUTP (ENZO Cy3-dUTP) was injected into the distal tip of the gonad of 20-24 hours post-L4 worms. Worms were dissected and immunostained 2.5 hours after injection.

### DNA damage sensitivity experiments

Young adult homozygous *fncm-1* animals were picked from the progeny of *fncm-1/hT2* parent animals. To assess for IR sensitivity, animals were treated with 0 and 50 Gy of γ-IR from a Cs^137^ source at a dose rate of 1.8 Gy/min. HU sensitivity was assessed by placing animals on seeded NGM plates containing 0, 3.5 and 5.5 mM HU for 12-16 hours. For interstrand crosslink sensitivity, animals were treated with 0 and 25µg/ml of Trioxsalen (trimethylpsoralen; Sigma) in M9 buffer with slow agitation in the dark for 30 minutes. Worms were exposed to 200 J/m^2^ of UVA. For all embryonic hatching assays, >36 animals were plated, 6 per plate, and hatching was monitored 60-72 hours after treatment as a readout of mitotic effects given how long it takes to proceed from the pre-meiotic region to egg laying (JARAMILLO-LAMBERT *et al.* 2010; KIM AND COLAIACOVO 2015a; KIM AND COLAIACOVO 2015b).

For larval arrest assays, L1 worms were plated on NGM plates with either 0 or 5.5 mM HU and incubated for 12-16 hours. The number of hatched worms and live adults were counted. Each damage condition was replicated at least twice in independent experiments as described in (KIM AND COLAIACOVO 2015a).

### Immunofluorescence and Western blot analysis

Whole mount preparations of dissected gonads, fixation and immunostaining procedures were carried out as described in (COLAIACOVO *et al.* 2003). Primary antibodies were used at the following dilutions: rabbit anti-SPR-5 (Santa Cruz sc-98749, 1:500), rabbit anti-SPR-5 ((NOTTKE *et al.* 2011), 1:1000 for western blot), rabbit anti-RAD-51 (SDI, 1:20000), rat anti-FCD-2 ((LEE *et al.* 2010), 1:300), rat anti-RPA-1 ((LEE *et al.* 2010), 1:200), rabbit anti-pCHK1 (Santa Cruz sc17922, 1:50), chicken anti-GFP (Abcam ab13970, 1:400), and mouse anti-H3K4me2 (Millipore CMA303, 1:200). Secondary antibodies used were: Cy3 anti-rabbit, FITC anti-rabbit, Cy3 anti-rat, Alexa 488 anti-chicken, and FITC anti-mouse (all from Jackson Immunochemicals), each at 1:250. Immunofluorescence images were collected at 0.2µm intervals with an IX-70 microscope (Olympus) and a CoolSNAP HQ CCD camera (Roper Scientific) controlled by the DeltaVision system (Applied Precision). Images were subjected to deconvolution by using the SoftWoRx 3.3.6 software (Applied Precision).

For western blot analysis, age-matched 24 hours post-L4 young adult worms were washed off of plates with M9 buffer. 6x SDS buffer was added to the worm pellets, which were then flash frozen in liquid nitrogen and boiled before equal amounts of samples were loaded on gels for SDS-PAGE separation.

### Co-localization analysis

The co-localization tool in Softworx from Applied Precision was employed for co-localization analysis (ADLER AND PARMRYD 2010).

### Mass spectrometry analysis

Pellets of age-matched 24 hours post-L4 young adult worms (wild type or *spr-5*::GFP::HA) were flash-frozen in lysis buffer (50mM HEPES pH 7.4, 1mM EGTA, 3mM MgCl2, 300mM KCl, 10% glycerol, 1% NP-40 with protease inhibitors (Roche 11836153001) using liquid nitrogen and then ground to a fine powder with a mortar and pestle. Lysis buffer was added to the thawed worms and samples were sonicated for 30 cycles of 20 seconds each. The soluble fraction of the lysate was applied to a 0.45µm filter and applied to either anti-HA beads (Sigma E6779) or GFP-Trap (Chromotek gta-20) that were incubated at 4°C overnight. After 3 washes with lysis buffer lacking NP-40, the bound proteins were eluted with either 1mg/ml HA peptide (Sigma I2149) or 0.1M glycine and precipitated using the Proteo Extract Protein Precipitation Kit (Calbiochem 539180). The dry pellet was submitted to the Taplin Mass Spectrometry Facility (Harvard Medical School) for analysis. The wild type sample was used as a negative control to remove false positive hits.

### Co-immunoprecipitation

Co-immunoprecipitations were performed with worm lysates from FNCM-1 tagged animals (*rj45[fncm-1::GFP∷3xFLAG]).* Lysis buffer was added to the worm lysates and they were sonicated for 30 cycles of 20 seconds each. The soluble fraction of the lysates was applied to anti-flag M2 magnetic beads (Sigma) that were incubated at 4°C overnight. Interacting proteins were eluted with glycine buffer (pH 2). Eluates were used for Western blot analysis to confirm the interaction of SPR-5 and FNCM-1 proteins.

### Statistical analysis

Statistical comparisons between mutants and control worms were carried out using the two-tailed Mann-Whitney test with a 95% confidence interval. *p<0.05.

## RESULTS

### Mass spectrometry and co-immunoprecipitation analyses reveal that SPR-5 interacts with FNCM-1

The histone demethylase SPR-5 in *C. elegans* as well as its orthologs in humans have been reported to function in DSB repair (HUANG *et al.* 2007; KATZ *et al.* 2009; BLACK *et al.* 2010; NOTTKE *et al.* 2011; MOSAMMAPARAST *et al.* 2013; PENG *et al.* 2015). To better understand the roles played by SPR-5 in DNA damage repair throughout the germline, we applied a proteomic approach to search for its interacting partners. Specifically, we performed pull-downs with a CRISPR-Cas9 engineered transgenic line expressing the endogenous SPR-5 tagged with GFP and HA *(spr-5:*:GFP∷HA), which did not display either the embryonic lethality or DSB sensitivity observed in *spr-5* null mutants (Supplemental Figure 1), followed by liquid chromatography-mass spectrometry (LC-MS) analysis. The Fanconi anemia (FA) FANCM homolog in *C. elegans*, FNCM-1, was identified in two independent samples utilizing this strategy, each processed with α-HA and α-GFP antibodies (Table 1). The SPR-5 and FNCM-1 interaction was not detected in control worms with untagged SPR-5 (Table 1) suggesting that SPR-5’s interaction with FANCM/FNCM-1 is specific. Proteins previously shown to interact with SPR-5, such as SPR-1, the ortholog of human co-repressor CoREST, and RCOR-1, an ortholog of human REST co-repressors 2 and 3 (RCOR2 and RCOR3) (JARRIAULT AND GREENWALD 2002; LEE *et al.* 2008), were also identified indicating that the pull-down followed by LC-MS worked efficiently.

**Table 1.** SPR-5 interacting proteins identified by liquid chromatography-mass spectrometry (LC-MS) analysis.

| Protein<br>Name | SPR-5::GFP::HA | Control |
| --- | --- | --- |
| RCOR-1 | 45 | Not detected |
| SPR-1 | 18 | Not detected |
| SPR-5 | 268 | Not detected |
| FNCM-1 | 3 | Not detected |

To further validate the interaction between SPR-5 and FNCM-1, we utilized a functional CRISPR-Cas9 engineered transgenic line expressing endogenous FNCM-1 tagged with GFP and FLAG *(fncm-1*∷GFP∷ FLAG; Supplemental figure 2) in co-immunoprecipitation experiments. We detected SPR-5 in pull-downs done from *fncm-1*∷GFP∷ FLAG worm lysates with an *a-* FLAG antibody, further supporting an SPR-5 and FNCM-1interaction *in vivo* (Figure 1A).

**Figure 1.**
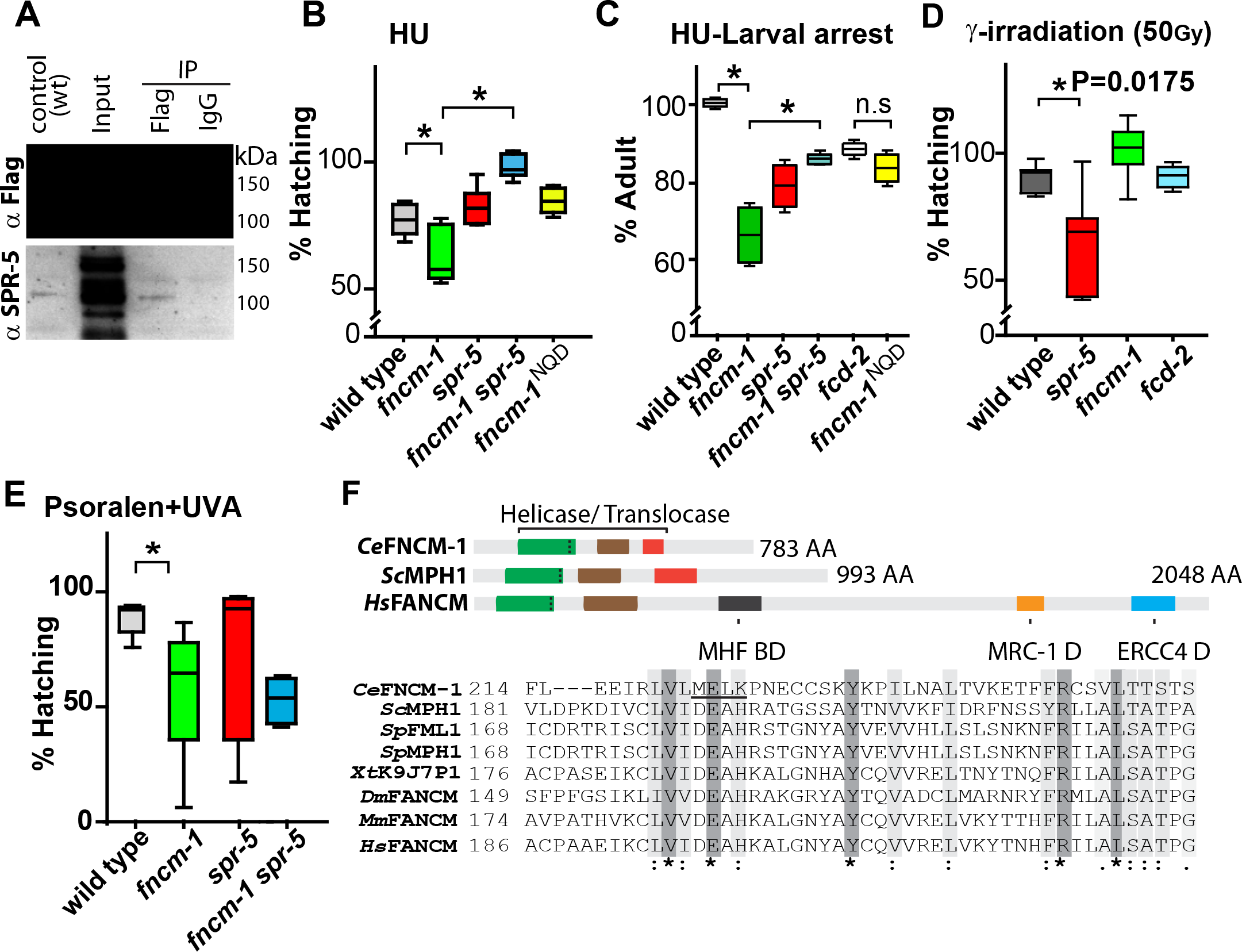
FANCM/CeFNCM-1 interacts with histone demethylase LSD1/CeSPR-5 and displays hydroxyurea-induced replication stress sensitivity that is suppressed in *spr-5* mutants. A. Western blots showing co-immunoprecipitation (co-IP) of FNCM-1 and SPR-5 from *fncm-1*∷GFP∷FLAG transgenic whole worm lysates with anti-FLAG and anti-SPR-5 antibodies, respectively. Input represents a concentrated whole worm lysate sample prepared for co-IP. A wild type (N2) worm lysate is shown as a control for anti-SPR-5 and anti-FLAG antibodies. IgG is used as a control for the IP. B. and C. Relative percentage of hatching and larval arrest for the indicated genotypes after treatment with 3.5mM and 5.5mM hydroxyurea (HU), respectively. Relative values are calculated against the absence of treatment. D. FNCM-1 and FCD-2 are not required for DSB repair. *fncm-1* and *fcd-2* mutants did not exhibit a decrease in embryonic viability (shown as % hatching) compared to wild type following exogenous DSB formation by γ-IR exposure (P=0.0530, 100% hatching for *fncm-1* and P=0.8357, 90% hatching for *fcd-2).* Asterisks indicate statistical significance. P values calculated by the two-tailed Mann-Whitney test, 95% C.I. E. Relative percentage of hatching for the indicated genotypes after treatment with and 25µg/ml of trimethylpsoralen-UVA. Asterisks indicate statistical significance; P values were calculated by the two-tailed Mann-Whitney test, 95% C.I. F. Representation of the helicase/translocase amino acid sequence conservation of *C. elegans* FNCM-1 and its homologs in *H. sapiens, M. musculus, D. melanogaster, X. tropicalis, S. pombe*, and *S. cerevisiae.* Alignment was performed using T-COFFEE and Pfam (http://pfam.sanger.ac.uk). Dark gray boxes (*) indicate amino acid identity and light gray boxes (:) indicate similarity. Three vertical dots inside the green boxes indicate the position of the represented amino acid sequence. The location of the MEK to NQD mutation in the *C. elegans* sequence is underlined (MELK).

### SPR-5 and FNCM-1 cooperate upon DNA replication fork arrest

Since our analysis supports the interaction of SPR-5 with FANCM/FNCM-1 and we previously demonstrated that SPR-5 is required for DSB repair (NOTTKE *et al.* 2011) we set out to gain insight into the link between SPR-5 and the FA pathway during DNA repair. To this end, we examined the sensitivity of *fncm-1* and *spr-5* null mutants to different types of DNA damage (LEE *et al.* 2010; NOTTKE *et al.* 2011). First, we found that *fncm-1* mutants displayed sensitivity to HU treatment, which results in replication arrest (Figure 1B and 1C). Specifically, only 61% of embryos hatched in *fncm-1* mutants compared to 75% for wild type (P=0.0367 by the two-tailed Mann-Whitney test, 95% C.I.) following an exposure to 3.5mM HU. Moreover, the HU sensitivity observed in *fncm-1* mutants was suppressed in *fncm-1 spr-5* double mutants (P=0.0006), while *spr-5* single mutants did not exhibit any sensitivity compared to wild type (P= 0.1120). Similarly, the increased larval arrest observed in *fncm-1* mutants following HU treatment was also suppressed in *fncm-1 spr-5* double mutants (Figure 1C). Taken together, these observations suggest that FNCM-1 and SPR-5 play a role in DNA repair following collapse of stalled replication forks.

Next we examined the DNA damage sensitivity of *spr-5* and FA pathway mutants to exogenous DSBs generated by γ-IR. A significant reduction in the levels of hatched embryos was observed in *spr-5* null mutants compared to wild type animals (Figure 1D, 61% and 89% respectively, at a dose of 50Gy; P=0.0175 by the two-tailed Mann-Whitney test, 95% C.I.). However, both *fncm-1* and *fcd-2* null mutants, which lack the FANCD2 homolog in worms, were not sensitive to exogenous DSBs (100% and 90% hatching, respectively) suggesting that the FA pathway is not involved in DSB repair.

Analysis of the sensitivity to DNA interstrand crosslink’s (ICLs) revealed that *spr-5* mutants were not sensitive to ICLs induced by psoralen-UVA in germline nuclei (Figure 1E). Specifically, 75% of embryos laid by *spr-5* mutants hatched compared to 94% in wild type (P=0.2307 by the two-tailed Mann-Whitney test, 95% C.I.). However, as expected given that FNCM-1 is required for ICL repair (COLLIS *et al.* 2006), *fncm-1* mutants exhibited significant sensitivity as shown by only 55% hatching (P=0.0087). *fncm-1 spr-5* double mutants did not alter the sensitivity observed in *fncm-1* single mutants (57%, P=0.9176) indicating that SPR-5 does not play a role in ICL repair in germline nuclei (Figure 1E). Altogether, these observations suggest that the FA pathway may not be involved in DSB repair in conjunction with SPR-5 and that SPR-5 does not participate in ICL repair along with the FA pathway, but that instead their interaction is necessary upon DNA replication fork arrest.

### A potential helicase/translocase domain in FNCM-1 is important for somatic repair

The FANCM *C. elegans* homolog FNCM-1 contains well-conserved helicase/translocase domains also present from budding yeast to humans (Figure 1F). We generated a helicase/translocase dead mutant by CRISPR-Cas9 engineering based on helicase/translocase dead mutants produced in the MPH1 gene in yeast (SCHELLER *et al.* 2000) that contains the following amino acid changes: M to N, E to Q, and K to D at positions 247, 248, and 250. Interestingly, the *fncm-1^NQD^* mutant exhibited larval arrest levels similar to that observed in *fcd-2* null mutants (Figure 1C and 1F, P=0.0247; 84% adults for NQD and 100% for wild type, values are normalized against untreated controls), suggesting that the potential helicase/translocase domain (M^247^ E^248^ K^250^) is important for somatic repair. However, this helicase/translocase domain is not necessary for DNA repair upon replication fork arrest in the germline (Figure 1B, P=0.0017 and P=0.7206, compared to *fncm-1* and wild type respectively).

### FNCM-1 promotes replication fork progression and SPR-5 is required for the formation of single-stranded DNA regions induced by FNCM-1 deficiency

Since FANCM has been implicated in promoting S-phase progression (WHITBY 2010), we hypothesized that FNCM-1 might have a similar role. To address FNCM-1’s potential role in S-phase progression, we monitored the incorporation of a fluorescent nucleotide during S-phase by injecting Cyanine-3-dUTP in the *C. elegans* gonad. Although we did not observe overt differences in the overall length of the gonads in the mutants compared to wild type, we accounted for this possibility by assessing the relative distance of Cy3-labeled nuclei. We divided the distance of Cy3-labeled nuclei from the distal tip by the length of the specific gonad from distal tip to late pachytene. The relative distance between the Cy3-labeled nuclei and the distal tip was reduced significantly in *fncm-1* mutant germlines compared to wild type, suggesting a slowdown in the rate of S-phase progression in the *fncm-1* mutants (Figure 2, relative distance of 7.4 for *fncm-1* and 9.5 for wild type, P<0.0001). Consistent with the HU sensitivity assay, the slowdown in S-phase progression observed in *fncm-1* single mutants was suppressed in *fncm-1 spr-5* double mutants (P<0.0001). Furthermore, *fncm-1^NQD^* mutants also displayed a slowdown in S-phase progression, albeit not as severe as that observed for *fncm-1* null mutants, suggesting that the *fncm-1^NQD^* mutant is likely a hypomorphic allele (Figure 2, 7.4 for *fncm-1* and 8.7 for *fncm-1^NQD^*, P=0.0078. Relative distance of 9.5 for wild type and 8.7 for *fncm-1^NQD^,* P=0.3165). To further validate the Cy3 labeling results we examined the formation of single-stranded DNA regions as a result of replication blockage by assessing the presence of RPA-1 signal, which localizes to single-stranded DNA. RPA-1 signal was detected following treatment with 3.5mM HU in *fncm-1* mutants but not in either wild type or *spr-5* null mutants (Figure 3A). Moreover, the RPA-1 signal observed in *fncm-1* mutants was suppressed in the *fncm-1 spr-5* double mutants. Taken together, these observations suggest that FNCM-1 is required for replication fork progression upon DNA damage and that SPR-5 may be involved in the formation of the single-stranded DNA regions induced upon absence of FNCM-1 function.

**Figure 2.**
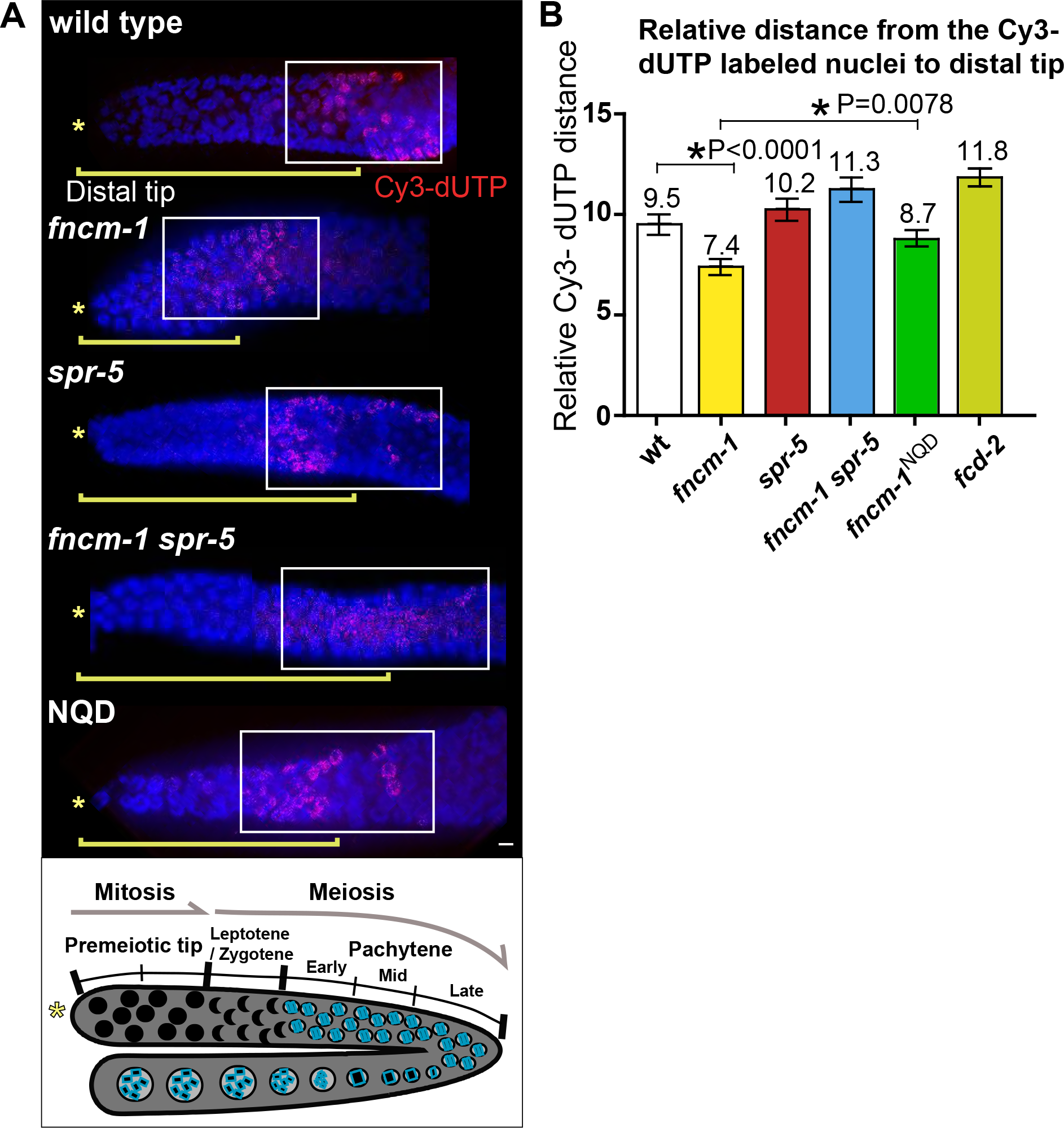
FNCM-1 is required for S-phase progression and impaired S-phase progression in *fncm-1* mutants is suppressed by lack of SPR-5. A. Cyanine-3-dUTP was injected into *C. elegans* gonads to monitor S-phase progression. Top, the distance between the Cy3-labeled nuclei and the distal tip (*) was measured for at least four gonads for each of the indicated genotypes. Bar, 2 µm. Bottom, diagram of the *C. elegans* germline indicating the mitotic (premeiotic tip) and meiotic stages represented in the top panel. B. Quantitation of the relative distance between Cy3-labeled nuclei and the distal tip in the germlines of the indicated genotypes. To account for potential variations in gonad size, the distance of Cy-3 labeled nuclei from the distal tip is divided by the length of the specific gonad from distal tip to late pachytene. Relative distance of Cy3-labeled nuclei = the distance of Cy3-labeled nuclei from the distal tip/the length of the specific gonad from distal tip to late pachytene x 100. At least four gonads were scored for each. Asterisks indicate statistical significance; P values calculated by the two-tailed Mann-Whitney test, 95% C.I.

**Figure 3.**
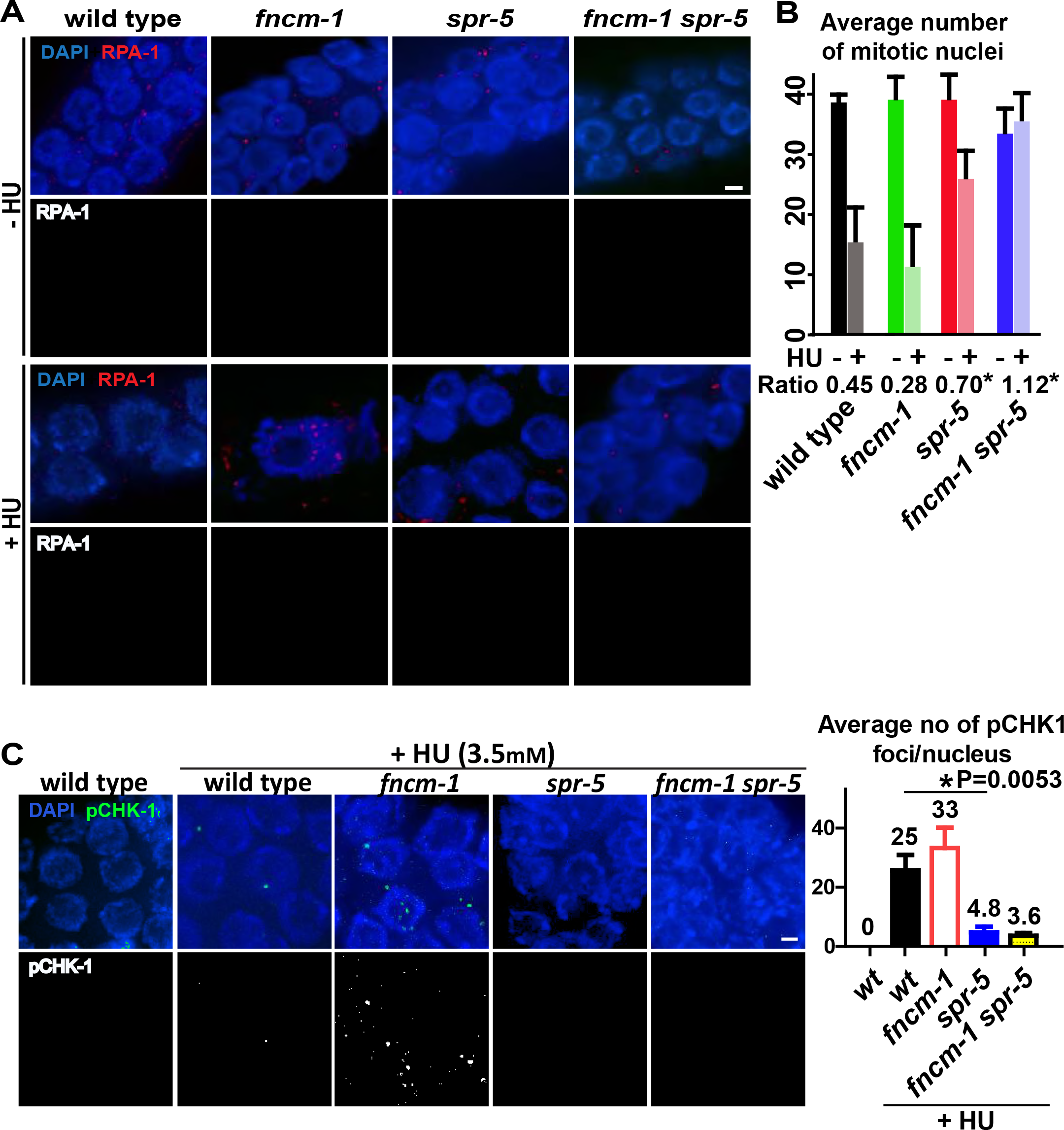
FNCM-1 promotes replication fork progression and SPR-5 is required for the S-phase checkpoint sensing the single-stranded DNA region formed upon lack of FNCM-1. A. Immunolocalization of single-stranded DNA binding protein RPA-1 upon 3.5mM HU treatment in the premeiotic tip for the indicated genotypes. Bar, 2 µm. B. Quantitation of the average number of mitotic nuclei within 40 µm in the premeiotic tip region of the germlines from the indicated genotypes. Ratio represents the number of nuclei observed following HU treatment (+ HU) divided by the number observed without treatment (-HU). Asterisks indicate statistical significance compared to wild type control. P=0.0422 for *spr-5,* P =0.0095 for *fncm-1 spr-5.* P values calculated by the two-tailed Mann-Whitney test, 95% C.I. C. S-phase DNA damage checkpoint activation is impaired in *spr-5* single and *fncm-1 spr-5* double mutants. Left, immunostaining for phospho CHK-1 (pCHK-1) on germline nuclei at the premeiotic tip following 3.5mM HU treatment. Bar, 2 µm. Right, quantitation of pCHK-1 foci. P values calculated by the two-tailed Mann-Whitney test, 95% C.I.

### S-phase DNA damage checkpoint activation is SPR-5-dependent

The DNA replication-dependent S-phase checkpoint is activated upon stress, such as HU treatment, DNA damage, and the presence of abnormal DNA structures, and results in S-phase arrest which is characterized by a premeiotic tip exhibiting enlarged nuclear diameters as well as a reduced number of nuclei in the *C. elegans* germline (BARTEK *et al.* 2004; GARCIA-MUSE AND BOULTON 2005; KIM AND COLAIACOVO 2014). Since the *spr-5* null mutation suppressed the single-stranded DNA formed in *fncm-1* we examined whether SPR-5 is required for the activation of the S-phase DNA damage checkpoint.

The ratio of mitotic nuclei (+HU/-HU) was not significantly changed in *fncm-1* mutants compared to wild type suggesting that the S-phase checkpoint is intact (Figure 3B, 0.45 and 0.28, respectively). However, a significant increase in the number of nuclei was observed in both *spr-5* single (0.70) and *fncm-1 spr-5* (1.126) double mutants compared to wild type (P=0.0422 and P=0.0095, respectively), indicating that SPR-5 is required for the S-phase DNA damage checkpoint and that lack of SPR-5, which causes accumulation of active chromatin (KATZ *et al.* 2009; NOTTKE *et al.* 2011), circumvents proper activation of the S-phase checkpoint.

Single-stranded DNA formed at a stalled replication fork is recognized by RPA and this triggers ATR kinase activation, which results in S phase checkpoint activation by phosphorylating its downstream target checkpoint kinase 1 (Chk1) (CIMPRICH AND CORTEZ 2008). Consistent with our observations of an impaired S-phase checkpoint, such as the increased number of mitotic germline nuclei as well as suppressed detection of single-stranded DNA, we detected a decrease in the levels of phosphorylated CHK-1 (pCHK-1) in these nuclei in *spr-5* mutants compared to wild type upon 3.5mM HU treatment (Figure 3C, P=0.0053). Altogether, these data indicate that SPR-5 is required for S-phase DNA damage checkpoint activation.

### The localization of SPR-5 and the FA pathway components FCD-2, FAN-1, and FNCM-1 is altered in response to replication stress

Since our analysis links SPR-5 with the FA pathway via FNCM-1 as functioning at stalled replication forks, we examined the localization of SPR-5 and factors acting in the FA pathway by immunostaining. Consistent with our previous observation, SPR-5 shows a nuclear-associated pattern (NOTTKE *et al.* 2011). Interestingly, upon HU treatment, we observed an increase in both peri-chromosomal SPR-5 signal as well as bright foci on chromatin compared to untreated (-HU) control wild type (Figure 4A), suggesting a role for the histone demethylase at replication fork arrest during S-phase. We also observed a brighter and elevated number of FANCD2/FCD-as well as FAN1/FAN-1 chromatin-associated foci following HU treatment, which supports the function of the *C. elegans* Fanconi Anemia pathway at stalled DNA replication forks, analogous to recent reports in other species (Figure 4A; (LACHAUD *et al.* 2016; MICHL *et al.* 2016)). FNCM-1∷GFP∷FLAG signal was observed as a combination of foci associated with the DAPI-stained chromosomes as well as a diffuse haze throughout the germline, which was not detected in the control wild type (Figure 4B and Supplemental figure 2). FNCM-1::GFP::FLAG partly co-localized with FCD-2 in the absence of any stress (-HU). However, its localization was altered upon replication fork arrest (+HU), as shown by the reduction of the diffuse germline signal and increase in bright chromatin-associated foci, suggesting that FNCM-1 responds to replication stress similar to FCD-2 and FAN-1, consistent with reports in other species (Figure 4B; (XUE *et al.* 2008)). Furthermore, we observed a higher level of co-localization between FNCM-1 and FCD-2 in the mitotically dividing nuclei at the premeiotic tip and a reduction in the level of co-localization at the pachytene stage, which supports FNCM-1 and FCD-2’s role in replication fork arrest at the mitotic stage (Figure 4C).

**Figure 4.**
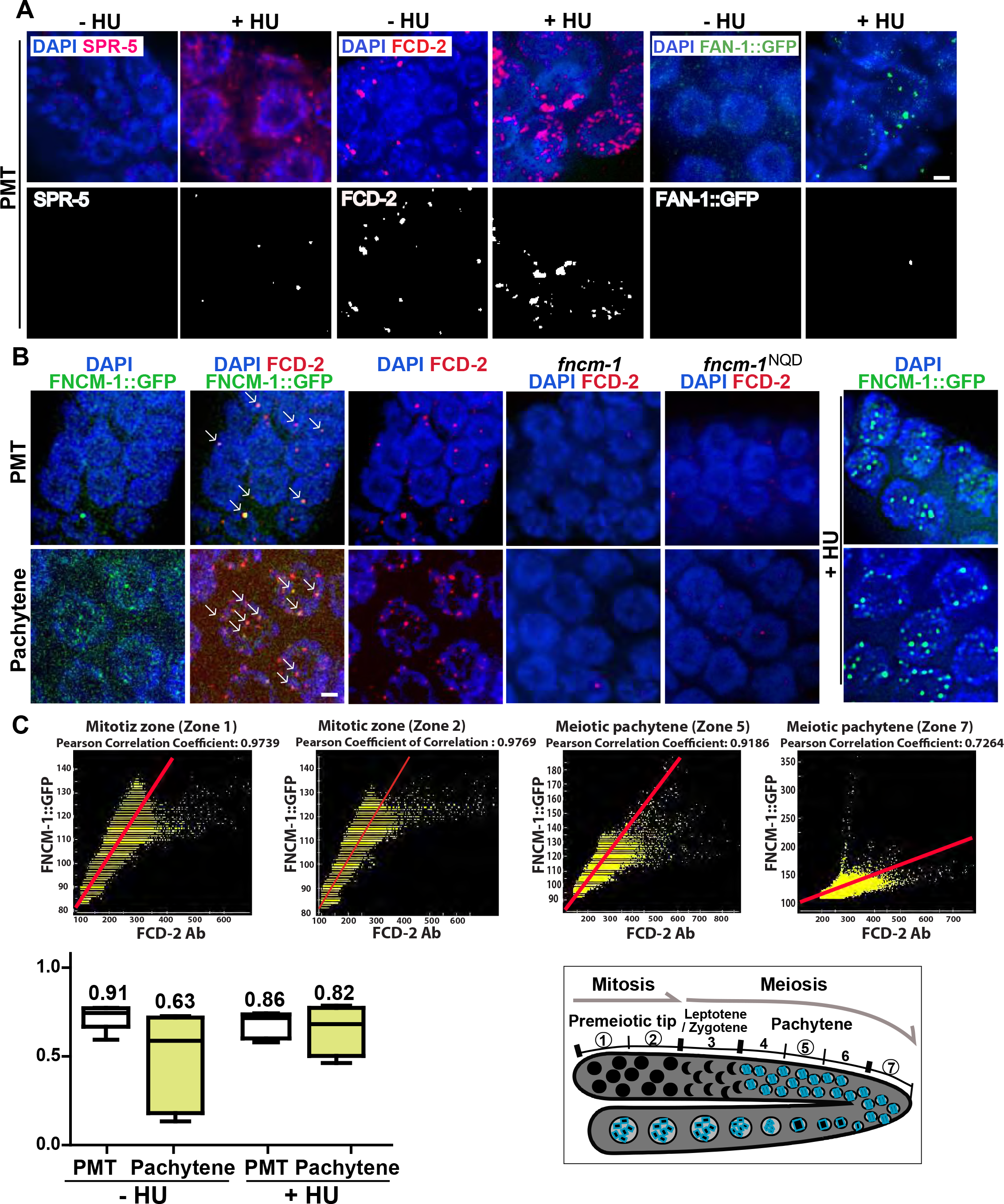
Proteins in the Fanconi anemia pathway and the histone demethylase SPR-5 display a dynamic localization upon HU treatment and co-localize. **A.** Immunolocalization of SPR-5, FCD-2 and FAN-1∷GFP (FAN-1∷GFP was detected with an anti-GFP antibody) upon 3.5mM HU treatment in the mitotically dividing germline nuclei (premeiotic tip, PMT). The localization pattern of SPR-5 and of the FA pathway components FCD-2 and FAN-1 is altered in response to replication stress (+HU). Bar, 2 µm. **B.** Immunolocalization of FNCM-1∷GFP and FCD-2 upon 3.5mM HU treatment in the premeiotic tip (PMT). FNCM-1 and FCD-2 co-localize on chromatin-associated foci (indicated by white arrows). Panel on the right shows that FNCM-1 re-localizes in response to replication stress changing from a more diffuse to a more focal localization. The dispersed FNCM-1∷GFP signal was not detected in control wild type (Supplemental figure 2). Bar, 2 µm. C. Top, graphs showing Pearson co-localization correlation coefficient values indicate higher co-localization levels between FCD-2 and FNCM-1∷GFP starting at the premeiotic tip (PMT) and slowly decreasing throughout meiosis (zones 1 and 2 = mitotic zone; zone 5 = mid-pachytene; zone 7 = late pachytene). Bottom left, mean numbers of Pearson co-localization correlation coefficient values between FCD-2 and FNCM-1∷GFP for both mitotic (PMT) and meiotic (pachytene) zones with or without HU exposure. n > 5 gonads. A value of indicates that the patterns are perfectly similar, every pixel that contains Cy3 (FCD-2, red) also contains GFP (FNCM-1∷GFP, green), while a value of −1 would mean that the patterns are perfectly opposite, every pixel that contains Cy3 does not contain GFP and vice versa. Bottom right, diagram of the *C. elegans* germline indicating the mitotic (zones 1 and 2) and meiotic stages (zones 5 and 7) represented in the top panel.

While FNCM-1 is known to be required for FCD-2 localization (Figure 4B, (COLLIS *et al.* 2006)), analysis of our helicase-dead *fncm-1^NQD^* mutant revealed a lack of FCD-2 localization, suggesting that the helicase/translocase domain is required for recruiting FCD-2 (Figure 4B). This is further supported by the observation that both *fcd-2* null and *fncm-1^NQD^* mutants displayed similar levels of larval arrest (Figure 1C) potentially due to the lack of FCD-2 localization in *fncm-1^NQD^* mutants mimicking *fcd-2* mutants. Taken together, these data support the idea that FNCM-1 responds to replication fork arrest and recruits downstream players FCD-2 and FAN-1 consistent with previous reports from other species. Also, we show for the first time that the helicase/translocase domain of FNCM-1is necessary for recruiting FCD-2.

### SPR-5 co-localizes with FNCM-1

Both SPR-5 and FNCM-1 localize from the premeiotic tip (PMT, mitotic zone) to pachytene (Figure 5A). We investigated whether SPR-5 and FNCM-1 co-localize on germline nuclei. However, since SPR-5 exhibits a dispersed localization, not limited to distinct foci, it is not possible to assess the co-localization of SPR-5 and FNCM-1 by scoring levels of superimposed foci. To circumvent this issue, we applied a Pearson correlation coefficient method (ADLER AND PARMRYD 2010). Consistent with their interaction by co-IP and LC-MS analysis, we found a high level of co-localization for FNCM-1 and SPR-5 (Figure 5B). Average Pearson correlation coefficient was 0.89 at the premeiotic tip and 0.80 at the pachytene stage in the germline. Interestingly, upon replication arrest following HU treatment, we found a high level of co-localization between SPR-5 and FNCM-1 persisting from the premeiotic tip to the pachytene stage, unlike in the control where this was progressively reduced. These observations support the idea that cooperation between the H3K4me2 histone demethylase and the FA pathway is reinforced to deal with replication fork blockage.

**Figure 5.**
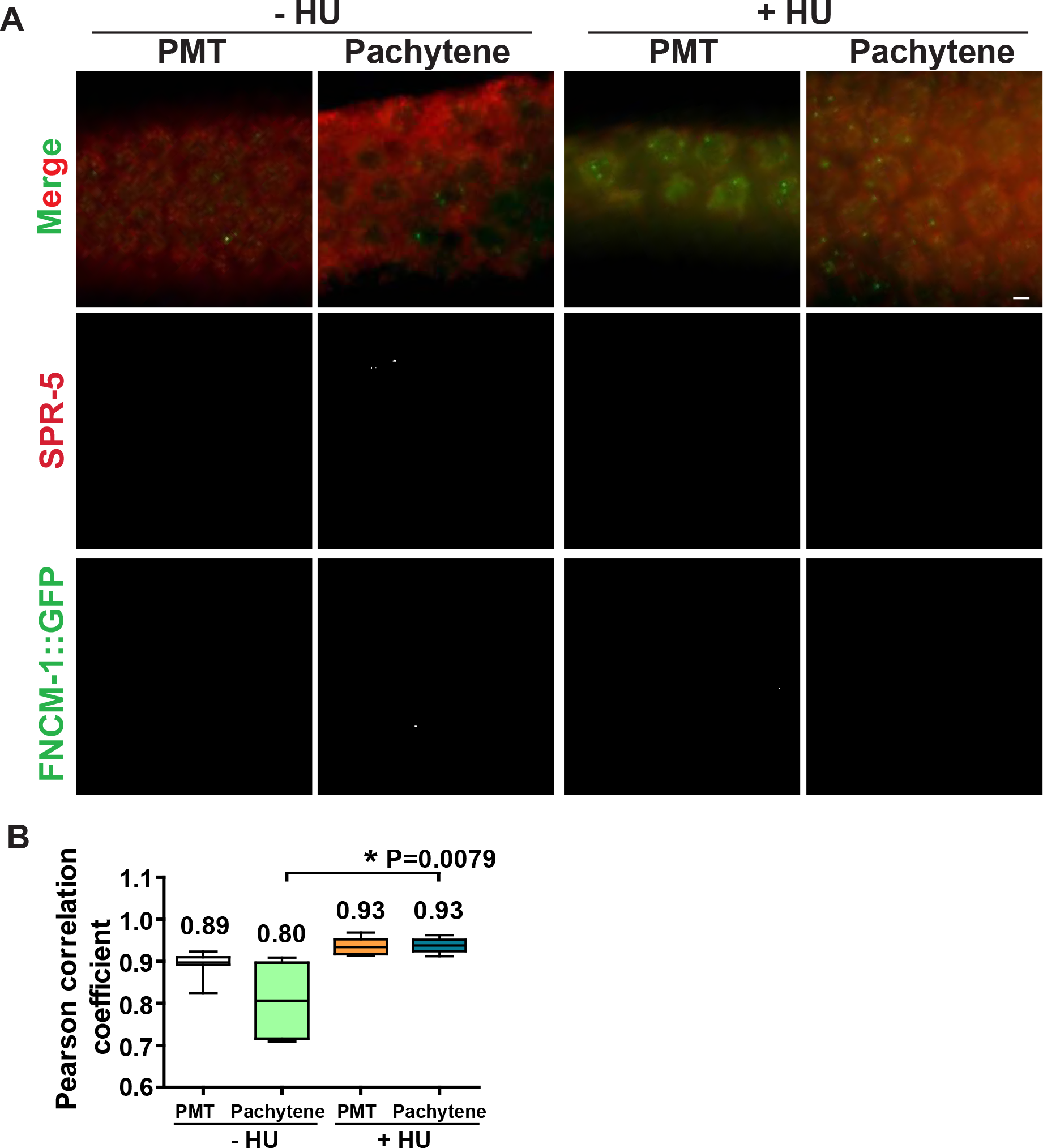
SPR-5 and FNCM-1 co-localization is extended upon replication fork stalling. **A.** Immunostaining of SPR-5 and FNCM-1∷GFP (endogenous signal) in nuclei at either the premeiotic tip (PMT) or at pachytene in the presence or the absence of 3.5mM HU treatment. Bar, 2 µm. **B.** Quantitation of Pearson co-localization correlation coefficient observed in A indicating that co-localization between SPR-5 and FNCM-1∷GFP extends into the pachytene stage upon replication fork stalling. 1 indicates a perfect positive linear relationship between variables. P=0.0079 for - and + HU treatment in the pachytene stage. P values calculated by the two-tailed Mann-Whitney test, 95% C.I.

### Two-way interaction of FANCM/FNCM-1 and LSD1/SPR-5: FNCM-1 and FCD-2 are necessary for maintaining proper H3K4 dimethylation levels

Since lack of LSD1/SPR-5 suppresses the HU sensitivity observed in *fncm-1* mutants, we next examined whether H3K4me2, which is regulated by the SPR-5 histone demethylase (KATZ *et al.* 2009; NOTTKE *et al.* 2011), was altered by the lack of FNCM-1. Surprisingly, we observed an increase in the levels of H3K4me2 in mutants lacking *fncm-1* suggesting a bidirectional functional interaction between SPR-5 and FNCM-1 (Figure 6A, numbers represent mean data from three independent experiments). Furthermore, H3K4me2 levels are also increased in *fcd-2* mutants, indicating that not only FANCM/FNCM-1 but also the FA pathway is necessary for maintaining histone demethylation together with LSD1/SPR-5. Although, we cannot rule out the possibility that the increase in H3K4me2 levels could be an indirect consequence resulting, for example, from alterations to cell cycle progression in the mutants.

**Figure 6.**
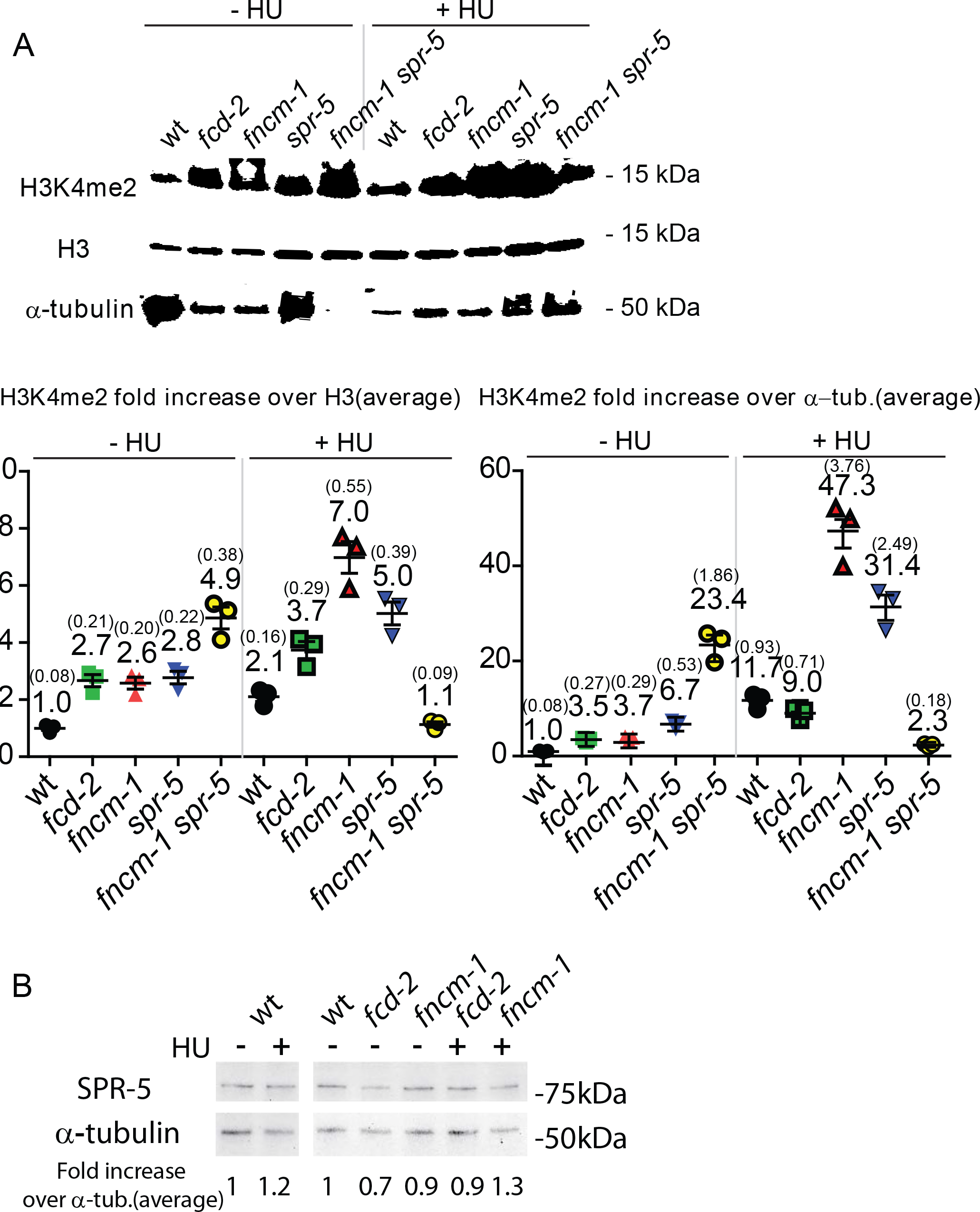
FNCM-1 and FCD-2 are necessary for maintaining H3K4 dimethylation levels. **A. Top,** Western blot analysis comparing H3K4me2 levels with histone H3 and alpha-tubulin antibodies for the indicated genotypes either in the absence or presence of HU (3.5mM). Bottom, Quantitation of H3K4me2 levels normalized against either histone H3 or alpha-tubulin. Signal intensity was measured with GelQuant.net. Numbers represent average for data from three independent experiments. SEM (standard error of mean) values are presented in parentheses. B. Western blot analysis comparing the levels of SPR-5 normalized against alpha-tubulin for the indicated genotypes upon absence or presence of 3.5mM HU treatment.

A previous study reported that human FANCD2 and FANCI are required for histone H3 exchange when cells are saturated with mitomycin C-induced DNA ICLs (SATO *et al.* 2012). Since defective H3 mobility possibly interferes with the accurate interpretation of H3K4 dimethylation levels, we normalized the H3K4me2 value to alpha-tubulin in addition to H3. Although we observed changes in the normalized level of H3K4me2, the overall conclusion from this analysis was not altered.

Since we observed an inverse correlation between the FA components and the levels of H3K4me2, we also examined the level of SPR-5 protein expression in *fncm-1*and *fcd-2* mutants in the absence or presence of HU exposure. However, the normalized expression level of SPR-5 against alpha-tubulin was not altered in wild type with or without HU exposure (Figure 6B). Also, FA mutants did not affect the level of SPR-5 expression regardless of HU exposure. These observations show that the level of H3K4me2 is not regulated by the level of expression of SPR-5 protein when replication forks stall. Taken together, our data support a two-way functional interaction between SPR-5 and the FA pathway in the germline: 1) in the activation of the S-phase DNA damage checkpoint in response to stalled replication forks, and 2) in the regulation of H3K4me2.

## DISCUSSION

Several studies have investigated the connections between epigenetic marks and DNA repair, however, the mechanisms by which epigenetic marks work in DNA repair remained unclear. Here, we show that the histone demethylase LSD1/CeSPR-5 interacts with the Fanconi Anemia FANCM/CeFNCM-1 protein based on biochemical, cytological and genetic analyses. LSD-1/CeSPR-5 is required for activation of the S-phase DNA damage checkpoint. Surprisingly, the FA pathway is required for H3K4me2 maintenance. Although a previous mouse study reported that FANCD2 modulates H3K4me2 at the sex chromosome, their analysis was confined to immunostaining (ALAVATTAM *et al.* 2016). With biochemical, cytological and genetic analyses, our study reveals that the FA pathway is necessary for epigenetic maintenance and sheds light on understanding the epigenetic mechanisms underlying Fanconi Anemia.

### The FA pathway responds to HU-induced replication fork arrest

The Fanconi Anemia pathway has been mainly studied in mitotically dividing cells but not in germline nuclei. In this study we identified a dynamic localization pattern for FNCM-1, FCD-2, FAN-1 and LSD-1/CeSPR-5 upon replication fork arrest induced by HU exposure (Figure 4A and 4B). In addition, co-localization, supported by an increased co-localization correlation coefficient, and co-IP results suggest that SPR-5 and FNCM-1 work together in response to replication fork arrest (Figures 1 and 5). Interestingly, *spr-5* mutants displayed DSB sensitivity but not the HU-induced replication fork sensitivity observed in *fncm-1* mutants (Figure 1D and Figure 1B, P=0.0175 and P=0.1720, respectively). However, a mild but significant reduction in larval arrest was observed (Figure 1C, P=0.0285, 100% for wild type and 79% for *spr-5*), suggesting a role for SPR-5 in mitotic cell division upon DNA replication stress. The interaction between these two proteins, as well as their altered localization upon HU stress, suggest that SPR-5 and FNCM-1 work together upon replication fork arrest.

### SPR-5 is necessary for activation of the DNA damage checkpoint

The S-phase checkpoint failure observed in *spr-5* mutants can be due to an impaired checkpoint signaling pathway *per se.* Suppression of the formation of a single-stranded DNA region in the *fncm-1 spr-5* double mutants (Figure 3A) suggests that SPR-5 may function in replication fork stalling/pause and that being deficient for SPR-5 prevents fork stalling, which then circumvents S-phase checkpoint activation. The defective checkpoint was observed at a lower (3.5mM) but not at a higher (5.5mM) dose of HU, suggesting that an alternative/redundant mechanism for S-phase checkpoint activation is triggered under severe replication stress conditions (Supplemental figure 3). It is worth noting that a similar role in checkpoint function was proposed in fission yeast for the Lsd1/2 histone demethylases which are indispensable for replication fork pause within the ribosomal DNA region (HOLMES *et al.* 2012).

### Fanconi Anemia components are required for proper H3K4me2 levels regardless of replication fork arrest

Surprisingly, FNCM-1 and FCD-2 were necessary to maintain proper H3K4me2 levels regardless of replication fork arrest (Figure 6). Since HU-induced replication arrest accumulates active chromatin marks during S-phase, the slowing down of S-phase observed in *fncm-1* mutants may result in H3K4me2 accumulation (Figure 2A, 3A and 6A, (ALPER *et al.* 2012)). However, this does not explain how FCD-2, which did not alter S-phase progression, is required for H3K4me2 with or without replication stress (Figure 2B and 6A). This suggests that, in addition to promoting the S-phase induced euchromatic state, the FA pathway may have an alternative role in maintaining histone methylation.

Although the FA pathway is connected to the regulation of histone demethylation regardless of the presence of stalled replication, a direct role for the FA pathway in histone demethylation became more evident when *fncm-1 spr-5* double mutants suppressed H3K4me2 upon HU arrest unlike either single mutant (Figure 6A). One possibility is that a defective checkpoint in *spr-5* somehow gains synergy in *fncm-1* mutants. Alternatively, a severe accumulation of dimethylation displayed in the double mutants may trigger/activate other histone demethylases. In fact, the LSD2 ortholog in *C. elegans, amx-1,* has been reported to be up-regulated over five-fold in *spr-5* mutants supporting this idea (KATZ *et al.* 2009; NOTTKE *et al.* 2011).

### The potential helicase domain (MEK) of FNCM-1 is necessary for recruiting FCD-2

Although *C. elegans* FNCM-1 displayed relatively less conservation of its DExD/H domain compared to other species, its flanking sequences are still well conserved (Figure 1F). Previous studies reported that the helicase domain of budding yeast Mph1, an ortholog of human FANCM, was required for mitotic crossover formation (PRAKASH *et al.* 2009). Interestingly, mutations in the potential helicase domain (MEK to NQD) of FNCM-1 resulted in loss of FCD-2 localization and a slowdown of S-phase progression (Figure 2). Moreover, it also led to larval arrest upon replication fork arrest comparable to that observed in *fcd-2* mutants, albeit not as severe as observed in *fncm-1* mutants (Figure 1C), which supports our observation that this domain in FNCM-1 is necessary to recruit the downstream FA pathway component FCD-2 (Figure 4B).

Fanconi Anemia (FA) is a rare genetic disorder but still the most frequent inherited instability syndrome, characterized by bone marrow failure, hypersensitivity to cross-linking agents and a high risk for acute myeloid leukemia, ataxia aelangiectasia, xeroderma pigmentosum, and Bloom, Werner, Nijmegen, Li-Fraumeni and Seckel syndromes (SCHROEDER 1982). Recent studies emphasize the role of Fanconi Anemia components in DNA replication arrest in addition to inter-strand crosslink repair (BLACKFORD *et al.* 2012; LACHAUD *et al.* 2016). Our findings that the FA pathway has a role in maintaining histone H3 K4 dimethylation regardless of replication stress supplies an important connection between DNA damage repair and epigenetic regulation (Figure 7). Furthermore, *fncm-1* mutants that are defective in recruiting FCD-2 will assist to define the precise contribution of the FA genes in this regulation.

**Figure 7.**
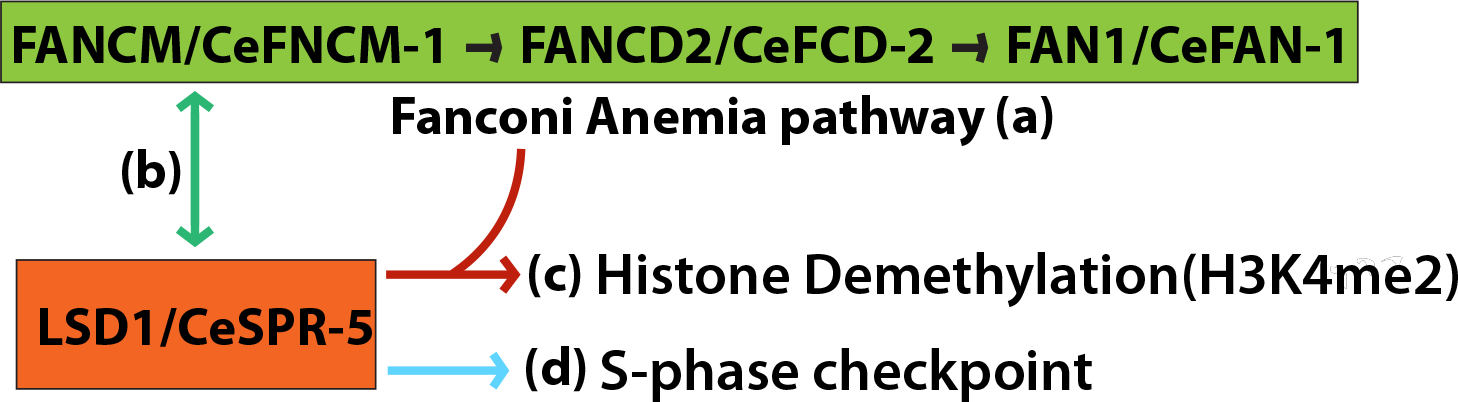
The FA pathway and SPR-5 cooperate in DNA repair and regulation of histone demethylation. **a.** *C. elegans* FNCM-1 is required for recruiting FCD-2 and its downstream nuclease FAN-1 in the germline. The potential helicase/translocase domain in FNCM-1 is necessary for this process. **b.** SPR-5 and FNCM-1 interact with each other and their co-localization in the germline is extended under conditions leading to stalled replication forks. **c.** FNCM-1 and FCD-2 are necessary for maintaining proper H3K4 dimethylation levels. **d.** SPR-5-dependent S-phase checkpoint activation is required in response to the single-stranded DNA region formed in the absence of FNCM-1 in germline nuclei.

## ACKNOWLEDGEMENTS

We thank Doris Lui for comments on the manuscript and members of Colaiácovo laboratory for discussions. We thank Dr. Koo for the RPA-and FCD-2 antibodies. This work was supported by a Ruth L. Kirschstein National Research Service award to SBS (F32GM100515) and a National Institutes of Health grant R01GM105853 to MPC.

---

**Supplemental figure 1.**
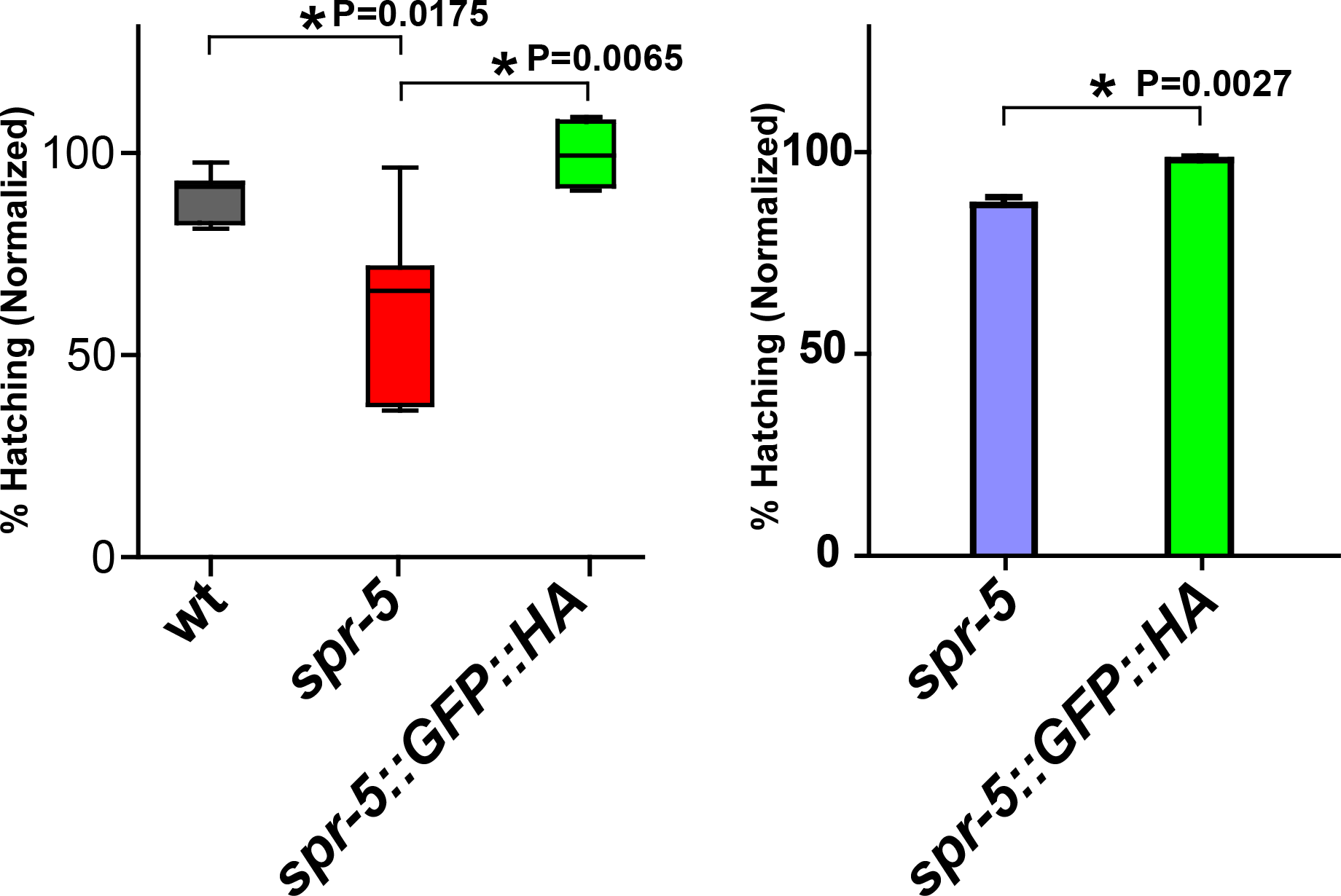
*spr-5*∷GFP∷HA animals generated by CRISPR-Cas9 do not display either sensitivity against exogenous DSBs induced by γ-IR (Left) or embryonic lethality (Right). Left, Normalized embryonic survival (shown as % hatching) was reduced in *spr-5* null mutants, but not in *spr-5:*:GFP∷HA worms, compared to control wild type animals following treatment with γ-irradiation (50 Gy), suggesting that insertion of the dual tags at the *spr-5* locus do not alter SPR-5 expression. 89% for wild type, 61% for *spr-5* and 98% for *spr-5*::GFP∷HA. P=0.0175 for *spr-5* compared to wild type by the two-tailed Mann-Whitney test, 95% C.I. P=0.0065 for *spr-5* compared to *spr-5*∷GFP∷HA. P=0.0931 for *spr-5*∷GFP∷HA worms compared to wild type. **Right**, *spr-5*∷GFP∷HA animals generated by CRISPR-Cas9 do not display embryonic lethality as in *spr-5* mutants. Embryonic survival was scored among the progeny of worms of the indicated genotypes. P=0.0027 for *spr-5* compared to *spr-5*∷GFP∷HA. Error bars represent standard error of the mean. n = at least 36 for each genotype.

**Supplemental figure 2.**
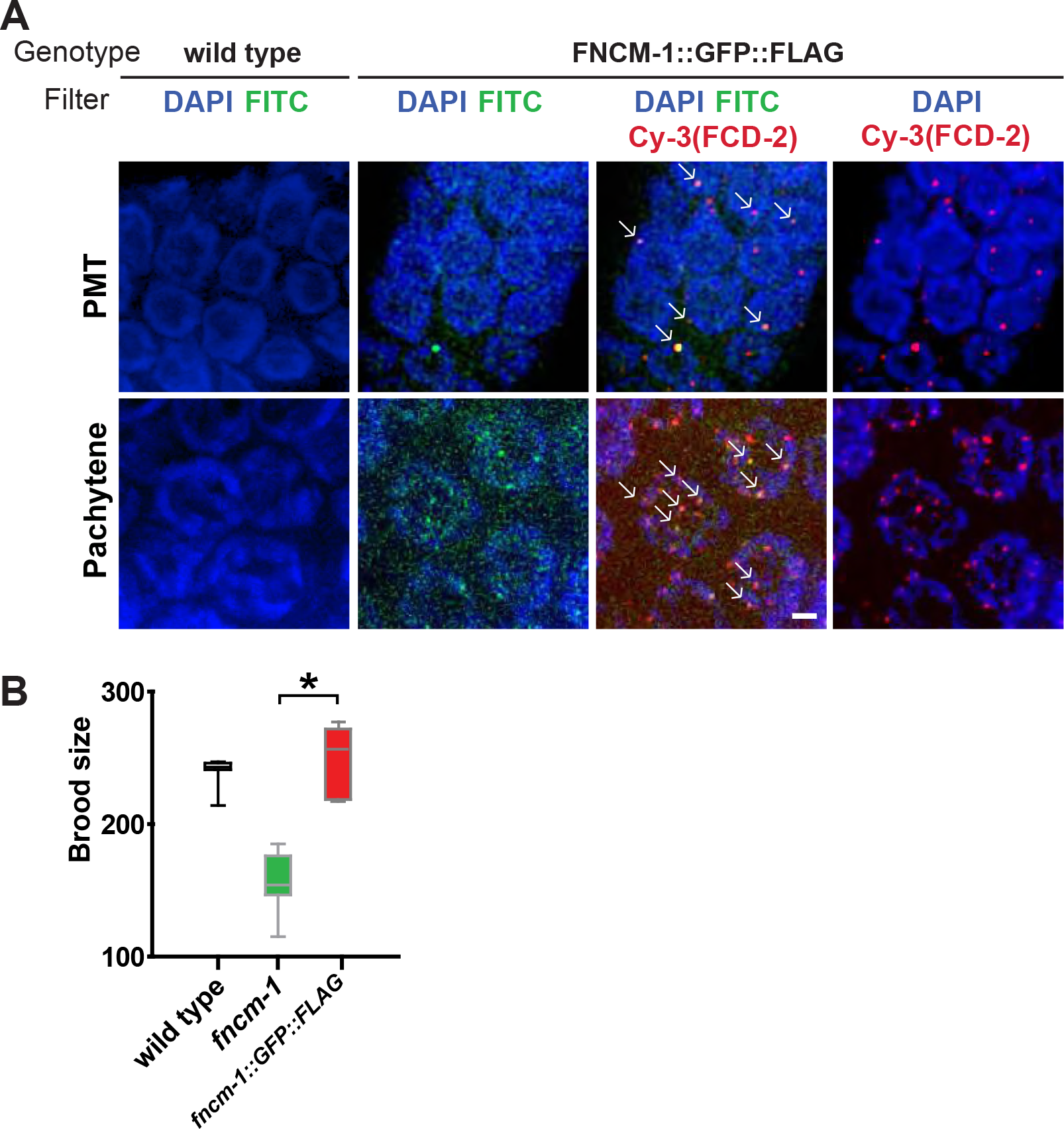
FNCM-1∷GFP∷FLAG co-localizes with FCD-2 and exhibits normal brood size. A. Endogenous FNCM-1∷GFP∷FLAG expression was detected as both foci on DAPI-stained chromatin as well as a dispersed signal observed throughout the germline, which was not detected in wild type. Localization of FNCM-1∷GFP∷FLAG clearly corresponded to FCD-2 indicated with white arrows. **B.** The *fcnm-1*∷GFP∷FLAG tag line displayed a brood size similar to wild type and significantly different (*) from *fncm-1* mutants suggesting that the *fncm-1*∷GFP∷FLAG transgenic line expressed wild type FNCM-1. Mean number of eggs laid per adult was 239 for wild type, 157 for *fncm-1*, and 249 for *fncm-1*∷GFP∷FLAG. P=0.0004 for *fncm-1* and *fncm-1*∷GFP∷FLAG. P=0.2234 for wild type and *fncm-1*∷GFP∷FLAG.

**Supplemental figure 3.**
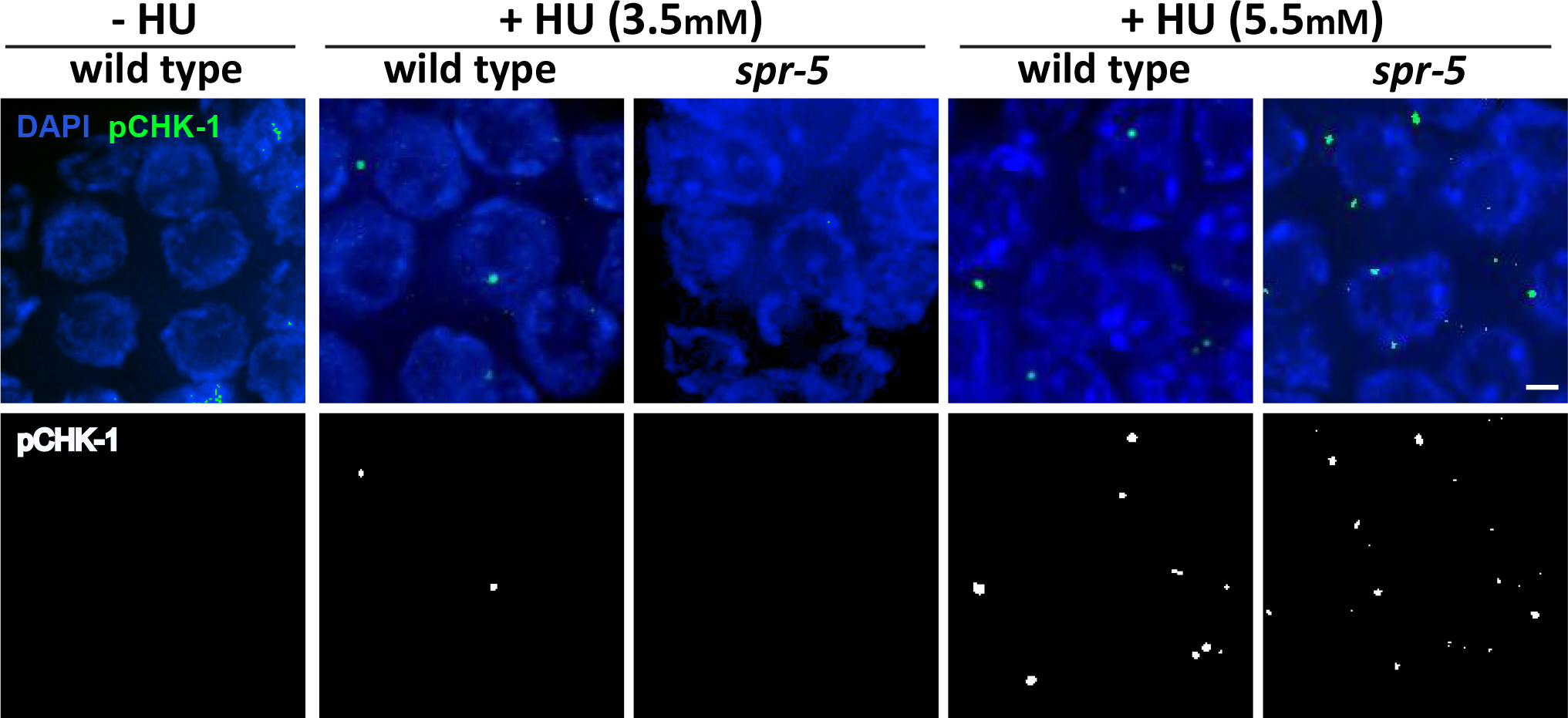
Defective checkpoint activation observed in *spr-5* mutants is HU dose-dependent. S-phase DNA damage checkpoint activation is impaired in *spr-5* mutants at a lower (3.5mM) but not at a higher (5.5mM) dose of HU. Immunostaining for phospho CHK-1 (pCHK-1) of mitotic nuclei (premeiotic tip) in the germline following either 3.5 or 5.5mM HU treatment. Bar, 2 µm.

